# Calling differential DNA methylation at cell-type resolution: an objective status-quo

**DOI:** 10.1101/822940

**Authors:** Han Jing, Shijie C. Zheng, Charles E. Breeze, Stephan Beck, Andrew E. Teschendorff

## Abstract

Due to cost and logistical reasons, Epigenome-Wide-Association Studies (EWAS) are normally performed in complex tissues, resulting in average DNA methylation profiles over potentially many different cell-types, which can obscure important cell-type specific associations with disease. Identifying the specific cell-types that are altered is a key hurdle for elucidating causal pathways to disease, and consequently statistical algorithms have recently emerged that aim to address this challenge. Comparisons between these algorithms are of great interest, yet here we find that the main comparative study so far was substantially biased and potentially misleading. By using this study as an example, we highlight some of the key issues that need to be considered to ensure that future assessments between methods are more objective.

---

DNA methylation (DNAm) is an epigenetic modification of DNA that is associated with gene regulation, and Epigenome-Wide Association Studies (EWAS) have identified significant numbers of genomic loci where DNAm is altered in association with disease risk factors and disease itself ^1, 2^. Most EWAS are performed in complex tissues made up of many underlying cell-types, which raises two immediate challenges for the interpretation of differentially methylated cytosines (DMCs) ^2–4^. First, there is the need to distinguish DMCs arising only because of changes in cell-type composition between cases and controls, from those that arise independently of these changes. For DMCs in the latter category, the second important challenge is to identify the specific cell-type(s) that drive the observed differential methylation. The importance of this task is clear: for instance, in the case of whole blood and a complex disease like Rheumatoid Arthritis, it can help reveal which immune cell subtype(s) is causally implicated ^5^. Or, in the context of epithelial cancer, it can reveal simultaneous DNAm changes in epithelial and fibroblast cells, and thus modulation of tumor-stroma paracrine signaling that could be critically important for cancer progression ^5^. While single-cell approaches provide, in theory, the best possible strategy to address this challenge, performing single-cell assays in large numbers of individuals, as required by an EWAS, is currently impossible, not least to generate reliable single-cell methylomes in spite of recent advances ^6^. Thus, there is an urgent need to perform cell-type deconvolution of DNA methylation profiles from bulk-tissue samples to reveal differentially methylated cell-types (DMCTs), that is DMCs at effectively cell-type resolution.

A number of statistical algorithms that aim to identify DMCTs have recently been proposed ^5 7–9^. One of these algorithms, called CellDMC, considers statistical interaction terms between phenotype and estimated cell-type fractions in a linear modelling framework, and is based on the intuition that differential methylation differences in a given cell-type would be larger in cases for which the affected cell-type is most abundant. Similar approaches that use statistical interactions or multiplicative effects between phenotype and cell-type have recently been proposed by Wu et al ^8^ and Luo et al ^9^. An entirely different algorithm called TCA, proposed by Rahmani and colleagues ^7^, attempts to first infer a data tensor defined over features, samples and cell-types, from which subsequent association analyses can be performed in each cell-type separately. This last study performed a direct comparison to CellDMC, concluding that TCA outperforms CellDMC. However, on inspecting the analysis and results of the TCA paper in more detail, we have identified numerous issues which, in our opinion, render the conclusions of the TCA paper invalid. According to our analysis and evaluation metrics, we find that CellDMC generally outperforms TCA, and that an objective comparison of the computational efficiency of the two methods clearly favors the application of CellDMC over TCA. Below, we describe some of the key issues, which in our opinion, have resulted in a severely biased analysis as reported in the TCA paper, in the hope that this may help to avoid such biases in future studies.

## Evaluation scenarios need to be precisely defined and comprehensive

Undoubtedly, evaluating a method that aims to identify DMCTs is a difficult task. Because for most EWAS datasets, the ground truth is completely unknown, simulation provides, in principle, the only means to objectively assess and compare competing methods. However, even for simulation there are many parameters and choices involved, which can greatly affect a method’s performance. In the context of identifying DMCTs, the number of parameters and scenarios is considerably larger than when identifying DMCs: for instance, it is not only the average effect size and number of samples that matter, but also the number of cell-types in the tissue, the distribution (notably average and variance) of relative cell-type proportions, whether changes are happening in one, two or any number of cell-types, in which direction alterations occurring in multiple cell-types happen (i.e. are they unidirectional in all affected cell-types, or can they be bidirectional), their relative effect sizes, and finally whether DMCTs occur at CpGs that are cell-type specific or not. Given this complexity, it is no surprise that the reporting of the simulation scenarios used in any given study is often highly ambiguous or lacks sufficient detail to ensure reproducibility.

Indeed, a concrete example is the TCA-study ^7^ where it is nowhere stated whether changes in underlying cell-types are uni or bi-directional. To demonstrate that this matters, we performed a comparative analysis of CellDMC to TCA in a scenario where two cell-types are changing and where at any given locus alterations in the two cell-types occur either in the same direction (unidirectional scenario) or in opposite direction (bidirectional scenario). Results for both methods in the unidirectional case are in line with those reported in the TCA paper, confirming that TCA displays improved sensitivity over CellDMC (Fig. 1A**, SI fig.S1**), although at the expense of a marginally lower specificity (Fig. 1B**, SI fig.S2**). However, for the bidirectional scenario we observed a widely different result, with TCA’s sensitivity exhibiting bounded behavior and dropping well below that of CellDMC (Fig. 1D**, SI fig.S3-S4**). Indeed, we have observed that TCA is only able to capture DMCTs that are directionally consistent with each other and which all change in a particular direction, failing to capture the DMCTs with opposite directionality (Fig. 1D, **SI fig.S3**), thus suggesting that TCA can’t detect bidirectional alterations. These results are robust to changes in the number of affected cell-types (**SI fig.S5-6**). Thus, we conclude that the data presented in the TCA paper is based on a simulation scenario where changes in multiple cell-types are unidirectional, and that TCA’s sensitivity is severely compromised in scenarios where bidirectional changes happen. Importantly, bidirectional changes in DNAm across multiple cell-types may not be that uncommon: for instance, in aging, bidirectional changes, although much less frequent than unidirectional ones, are also observed ^10^. Thus, it is clear that a range of different evaluation scenarios needs to be considered in order to perform an objective comparison between methods.

**Fig. 1:**
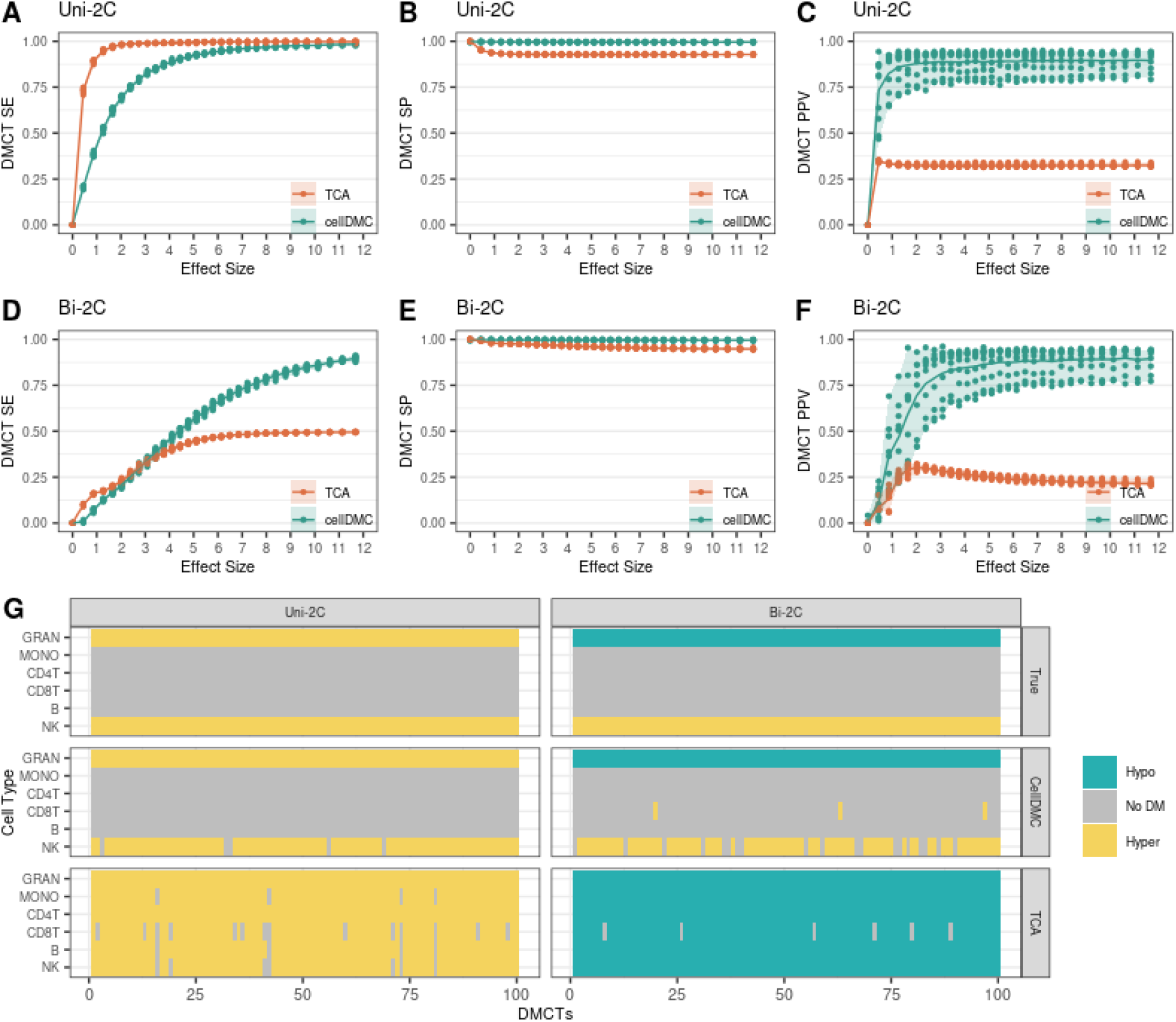
Performance metrics of CellDMC and TCA. **A)** Comparison of the sensitivity (SE) to detect DMCTs (y-axis) vs. effect size (x-axis) for the scenario where 2 of 6 blood cell subtypes are altered, with the loci in the two cell-types changing in the same direction (unidirectional 2 cell-type scenario, Uni-2C). Displayed are the average sensitivity over different cell-type pair combinations, for each of 10 Monte-Carlo runs. **B)** As A), but now for the specificity (SP) to detect DMCTs. **C)** As A), but now for the precision or positive predictive value (PPV). **D-F)** As A-C), but now for the case where the loci in the two altered cell-type exhibit opposite directionality of DNAm change (Bidirectional 2 cell-type scenario, Bi-2C). Solid lines represent the average over the 10 Monte-Carlo runs. **G)** Heatmaps displaying the significance and directionality of DNAm change for each blood cell subtype across 100 DMCTs for the two different scenarios Uni-2C and Bi-2C, with the top heatmap displaying the true patterns of change, and the middle and lower heatmaps displaying the predictions from CellDMC (middle) and TCA (lower). Displayed are the results for one typical Monte-Carlo run at an average effect size of 8.

## An objective evaluation requires multiple metrics

A second key issue is that comparisons of methods may often unjustifiably exclude important metrics ^11^. The TCA-paper illustrates this point. In the context of identifying DMCTs, obvious metrics to consider are the sensitivity (SE) or power to detect specific alterations, the specificity (SP) (related to the false positive rate (FPR) by SP=1-FPR), and the precision (or positive predictive value PPV). Indeed, the precision to correctly identify DMCTs is a critically important metric, yet the TCA-paper does not compare TCA to CellDMC in terms of this metric. We performed a comparison in terms of the PPV, finding that TCA’s precision is very low (Fig. 1C, Fig. 1F**, SI fig.S7-S9**). Indeed, in all of the scenarios analyzed here (many of which were also considered in the TCA paper), we find that TCA predicts DMCTs for cell-types that are not truly altered, even when effect sizes are large (Fig. 1G). This demonstrates that TCA suffers from an unacceptably high false discovery rate (FDR) or low precision.

Another important evaluation metric which was not considered in the TCA paper, and which is however specially relevant in the context of the TCA method, is the estimation of cell-type fractions themselves. Indeed, the TCA method involves an iterative procedure, in which initial cell-type fractions are re-estimated ^7^. A natural question to address therefore is whether the final cell-type fraction estimates agree better with the ground truth values, a question which is perfectly addressable not only within a simulation framework, but also in the context of real blood EWAS for which matched FACS blood cell counts are available. We have assessed this using both simulation and real blood DNAm datasets with matched FACS cell counts, concluding that TCA’s final cell-type fraction estimates are significantly worse than the initial ones (**SI fig.S10**). This contradicts the result shown in SuppFig.2 of the TCA paper, where in our opinion the authors used the wrong benchmark. For whole blood, reliable DNAm references exist ^12–14^ and yet the authors of the TCA paper initialize their TCA method with relatively poor cell-type fraction estimates, leading to the illusion that the final TCA estimates are good enough. Moreover, the authors of the TCA paper offer no assessment on real DNAm data with matched flow-cytometric cell counts, despite such data being available ^15^. Our analysis on both simulated and real DNAm data clearly demonstrates that TCA’s final cell-type fraction estimates are worse than those obtained using a gold-standard DNAm reference (**SI fig.S10**). It is therefore unclear to us how TCA could ever achieve an overall better performance.

## Evaluation metrics need to be meaningful

A third key problem with the TCA paper is the vague definition and even wrong use of evaluation metrics. In the context of identifying DMCTs, measures like sensitivity need to be defined very precisely, in order to avoid ambiguities and to ensure that the metric is meaningful. For instance, if we have a CpG that is truly hypermethylated in cell-type A, truly hypomethylated in cell-type B, and truly not differentially methylated in cell-type C, how do we treat it if a method predicts that it is hypermethylated in all 3 cell-types? While the prediction for cell-type A is correct, it is clearly incorrect for cell-types B and C. However, according to the definition of sensitivity as used in the TCA paper where it states “…*in the power simulations with effects in multiple cell types we considered a CpG to be associated with the phenotype if it had a significant association with at least one of the cell-types*”, the prediction for this CpG would have counted as being 100% correct across all cell-types. In other words, the sensitivity metric used by the authors of the TCA paper does not measure sensitivity to detect DMCTs. This example clearly illustrates the imperative need to define SE, SP and PPV metrics on a cell-type specific basis, and subsequently to average these values over all relevant cell-types in order to arrive at interpretable and meaningful metrics.

More worryingly, the TCA-study goes on to use their wrong sensitivity metric to make a comparison that is completely biased. Specifically, in Figure-2 of the TCA paper the authors use this wrong metric to compare the “sensitivity” of TCA and CellDMC to that of a standard linear regression method without interaction terms. To understand why their analysis is biased, we first note that a comparison of TCA or CellDMC to the standard linear method in terms of the sensitivity to detect DMCTs is completely meaningless, because by definition, a linear method without interaction terms can only identify DMCs (i.e whether a CpG is differentially methylated without regard of which cell-types have changed), and not DMCTs. On the other hand, a comparison of TCA or CellDMC to the standard linear model based on the author’s flawed sensitivity metric (a metric designed to quantify the power of detecting DMCs, not DMCTs) is biased and meaningless because TCA and CellDMC are designed to detect DMCTs. In other words, it defeats the whole purpose of presenting methods like TCA or CellDMC, if subsequently we don’t evaluate them in terms of their sensitivity to detect DMCTs, which is the task they were designed for. Indeed, the use of the wrong sensitivity metric explains why in Figure-2 of the TCA paper, CellDMC does not appear to outperform the standard linear regression model, a result which not only contradicts findings from several other studies ^5, 8^, but which is also statistically incorrect, because the additional interaction terms used in the CellDMC model *must* in general lead to an improvement over the standard model.

## Computational efficiency needs to be considered

When comparing computational algorithms a key evaluation metric is computational efficiency. This is particularly pertinent when the task at hand is computationally demanding. Because TCA infers a data tensor where the genomic feature (CpG) dimension is fairly large, it is a computationally very demanding task. In contrast, CellDMC scales linearly with the number of CpGs and does not take much longer to run than a standard linear regression model. Indeed, we performed a detailed comparison of the two methods in terms of their runtimes, finding that CellDMC is much more efficient (Fig. 2). For instance, running TCA on a typical EWAS study would take a day or more on a standard workstation, whilst CellDMC would complete the task in about 10 to 20 minutes (Fig. 2**, Table.S1**), which represents an approximate 100-fold improvement over TCA. Remarkably, the TCA-paper never compares methods in terms of their computational efficiency, nor is it ever mentioned that TCA requires such fairly long runtimes. The fact that this key limitation is never acknowledged in the manuscript further attests to their biased reporting.

**Fig. 2:**
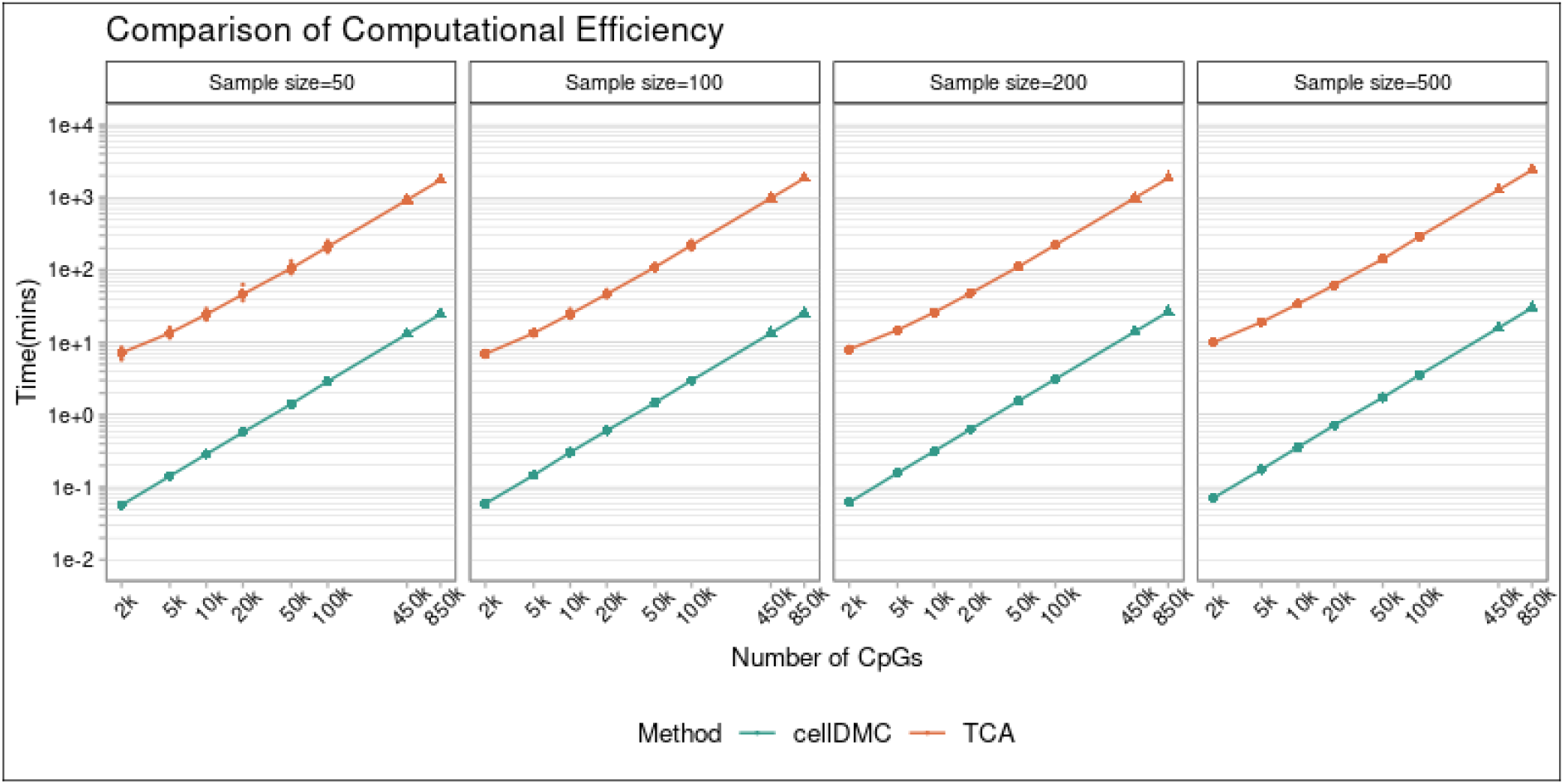
Runtimes of CellDMC and TCA. Runtimes (in minutes, shown in a log-base-10 scale) of CellDMC and TCA for 4 different sample sizes and different numbers of CpGs (x-axis, also shown on a log-base-10 scale). Runtimes are quoted for a standard workstation and by running the code on one core.

In summary, although previous articles ^11, 16 17^ have provided general guidelines to ensure more objective reporting of results, the TCA paper does not follow most of these guidelines, illustrating that overoptimistic or biased reporting persists. In the context of EWAS, identification of DMCTs at high sensitivity and precision is imperative to maximize the value of such studies. While TCA does exhibit improved sensitivity over CellDMC in specific scenarios, this comes at the expense of an unacceptably low precision or high false discovery rate. Our objective assessment demonstrates that overall, CellDMC achieves reasonably high sensitivity, specificity and precision over a wider range of different biological scenarios, and at a much lower computational cost compared to TCA.

## Declarations

### Author Contributions

Statistical and computational analyses were done by JH with help from SZ. CEB and SB contributed valuable feedback. Study was conceived and designed by AET. Manuscript was written by AET.

### Ethics approval and consent to participate

Not applicable, as this study only analyses existing publicly available data.

