## Supplementary Information for "Calling differential DNA methylation at cell-type resolution: an objective status-quo"

This document contains **Supplementary Tables, Supplementary Figures** and their legends, and **Supplementary Methods**.

### Supplementary Tables

| Time(min) | n = 200 | | n = 500 | |
|---|---|---|---|---|
| | ***cellDMC*** | ***TCA*** | ***cellDMC*** | ***TCA*** |
| **450k** | 15 | 1048 | 17 | 1356 |
| **EPIC** | 27 | 1875 | 30 | 2427 |
| **WGBS** | 62 | 4356 | 71 | 5637 |

**Table.S1: Computational Efficiency of cellDMC vs TCA.** We provide the running time (in minutes) for cellDMC and TCA for typical DNAm measurement technologies, which differ in the number of CpGs measured: approximately 480,000 CpGs for 450k, approximately 850,000 CpGs for EPIC and for WGBS we assumed 2.5 million CpGs.

### Supplementary Figures

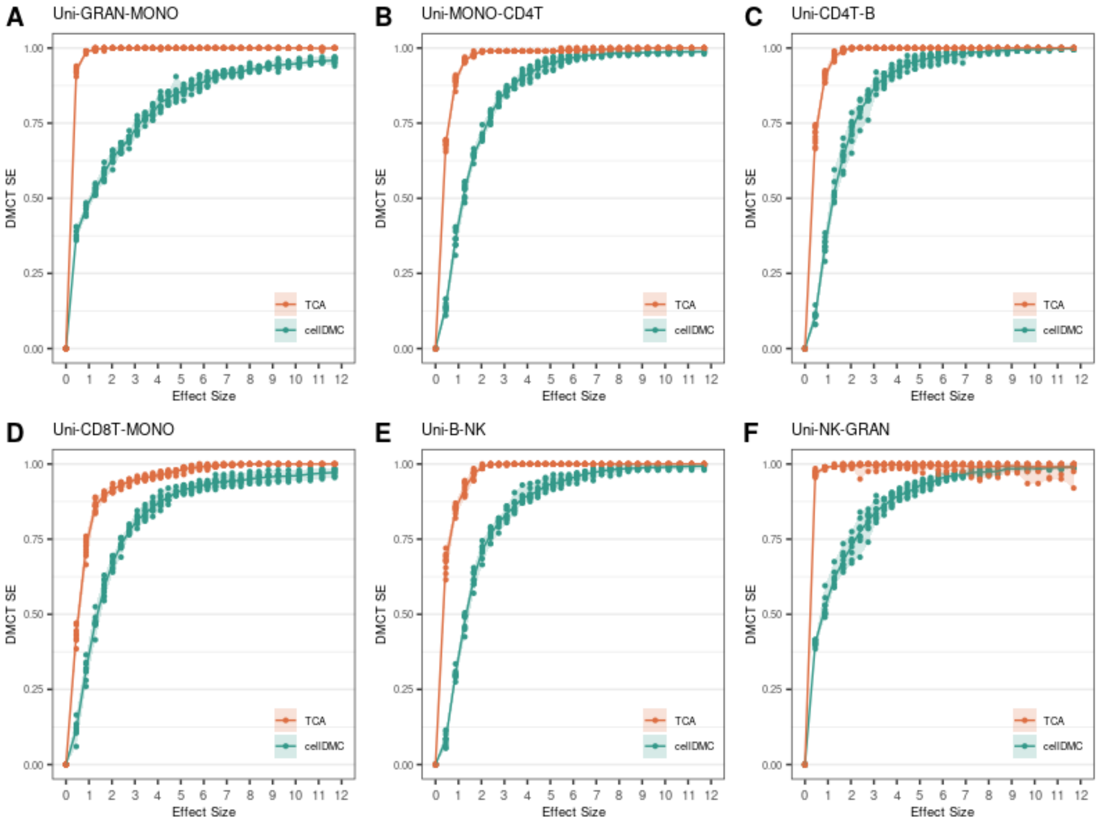


**Fig.S1: Comparison of the sensitivity (SE) in the unidirectional two altered cell-type scenario.** Plots of the sensitivity (SE) to detect DMCTs of CellDMC and TCA vs. the effect size for the scenario of unidirectional differentially methylation in 2 immune cell subtypes, as indicated above each plot. The mixtures are made of a total of 6 immune cell subtypes. Results are for 10 Monte Carlo runs, where in each run 200 samples were simulated (100 controls and 100 cases). The solid lines show the mean sensitivity change over the 10 Monte-Carlo runs.

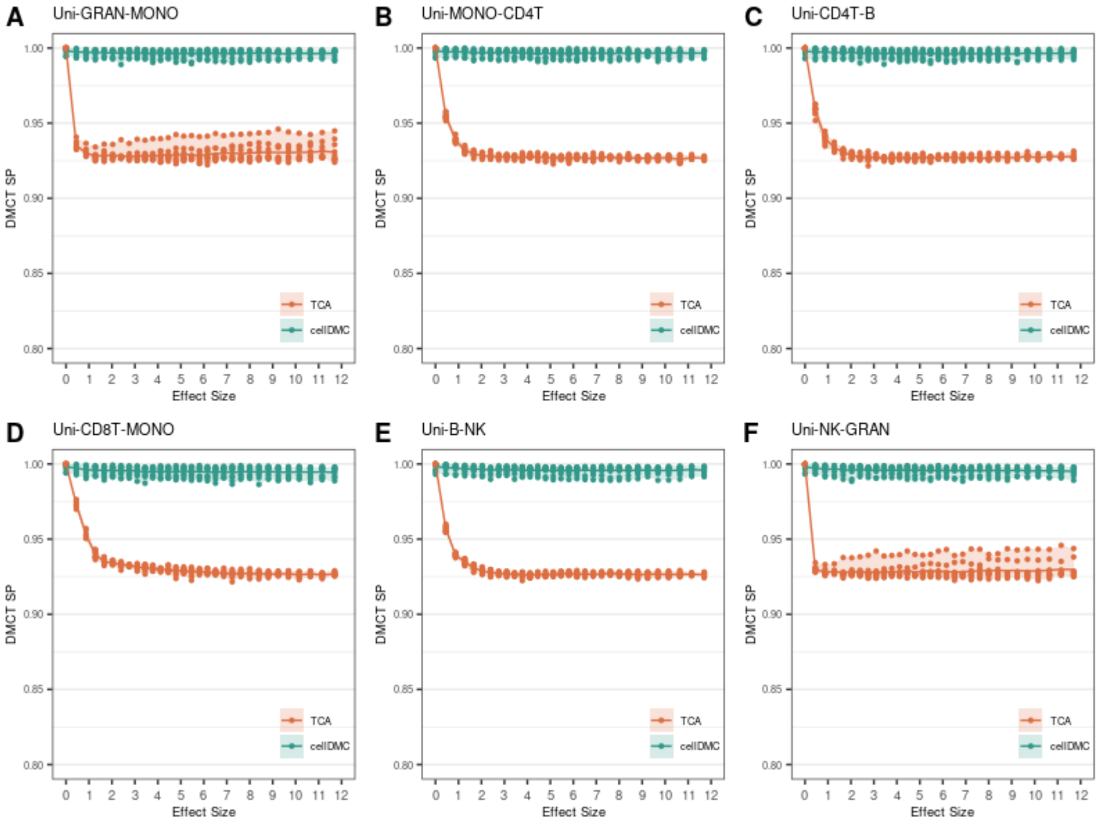


**Fig.S2: Comparison of the specificity (SP) in the unidirectional two altered cell-type scenario.** Plots of the specificity (SP) to detect DMCTs of CellDMC and TCA vs. the effect size for the scenario of unidirectional differentially methylation in 2 immune cell subtypes, as indicated above each plot. The mixtures are made of a total of 6 immune cell subtypes. Results are for 10 Monte Carlo runs, where in each run 200 samples were simulated (100 controls and 100 cases). The solid lines show the mean sensitivity change over the 10 Monte-Carlo runs

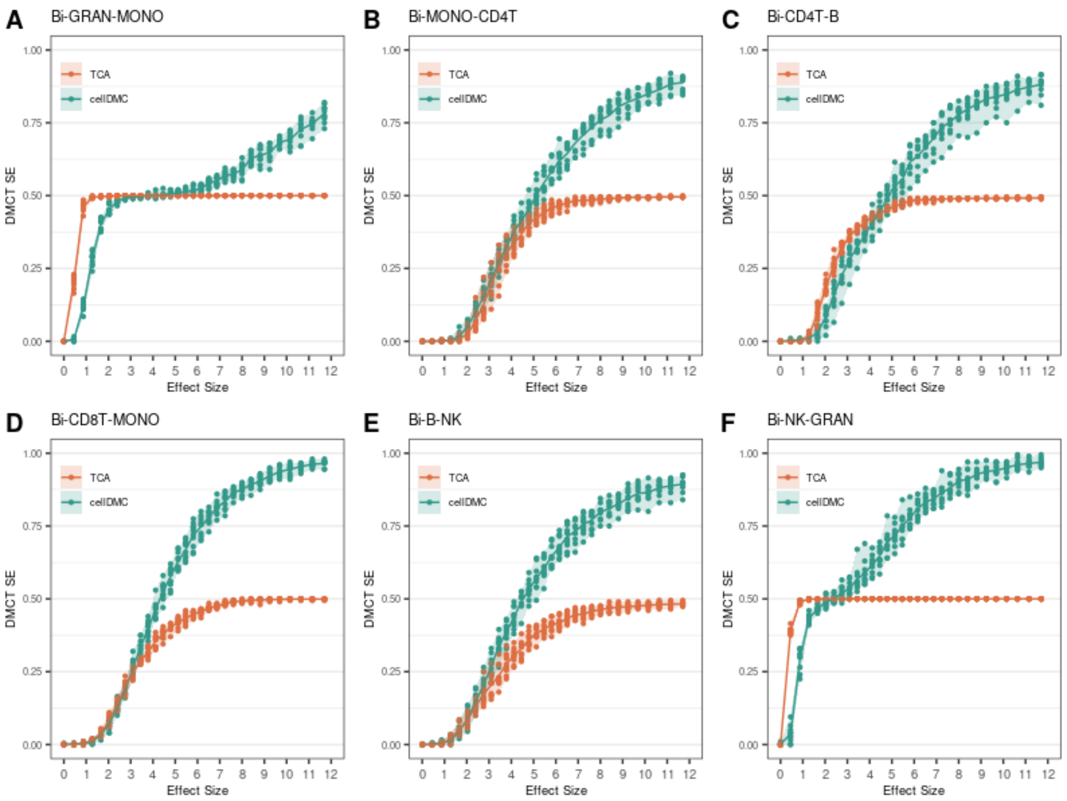


**Fig.S3: Comparison of the sensitivity (SE) in the bidirectional two altered cell-type scenario.** Plots of the sensitivity (SE) to detect DMCTs of CellDMC and TCA vs. the effect size for the scenario of bidirectional differentially methylation in 2 immune cell subtypes, as indicated above each plot. The mixtures are made up of a total of 6 immune cell subtypes. Results are for 10 Monte Carlo runs, where in each run 200 samples were simulated (100 controls and 100 cases). The solid lines show the mean sensitivity change over the 10 Monte-Carlo runs.

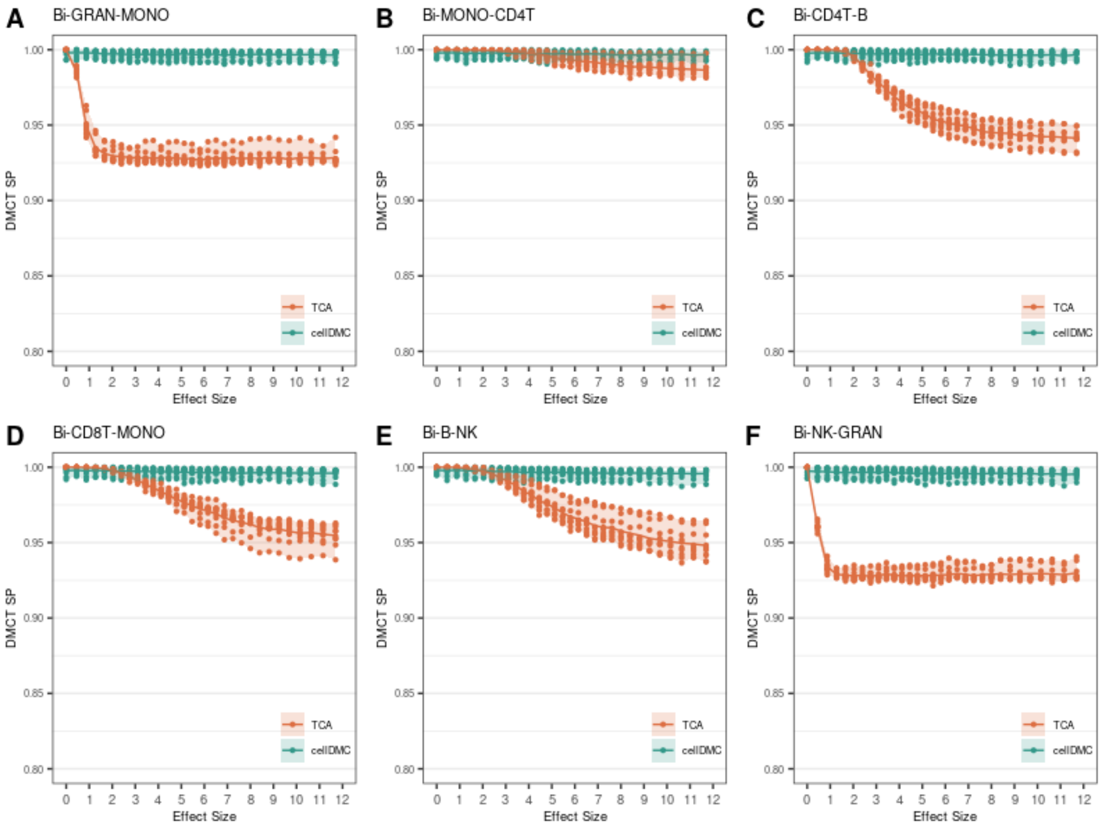


**Fig.S4: Comparison of the specificity (SP) in the bidirectional two altered cell-type scenario.** Plots of the specificity (SP) to detect DMCTs of CellDMC and TCA vs. the effect size for the scenario of bidirectional differentially methylation in 2 immune cell subtypes, as indicated above each plot. The mixtures are made up of a total of 6 immune cell subtypes. Results are for 10 Monte Carlo runs, where in each run 200 samples were simulated (100 controls and 100 cases). The solid lines show the mean sensitivity change over the 10 Monte-Carlo runs.

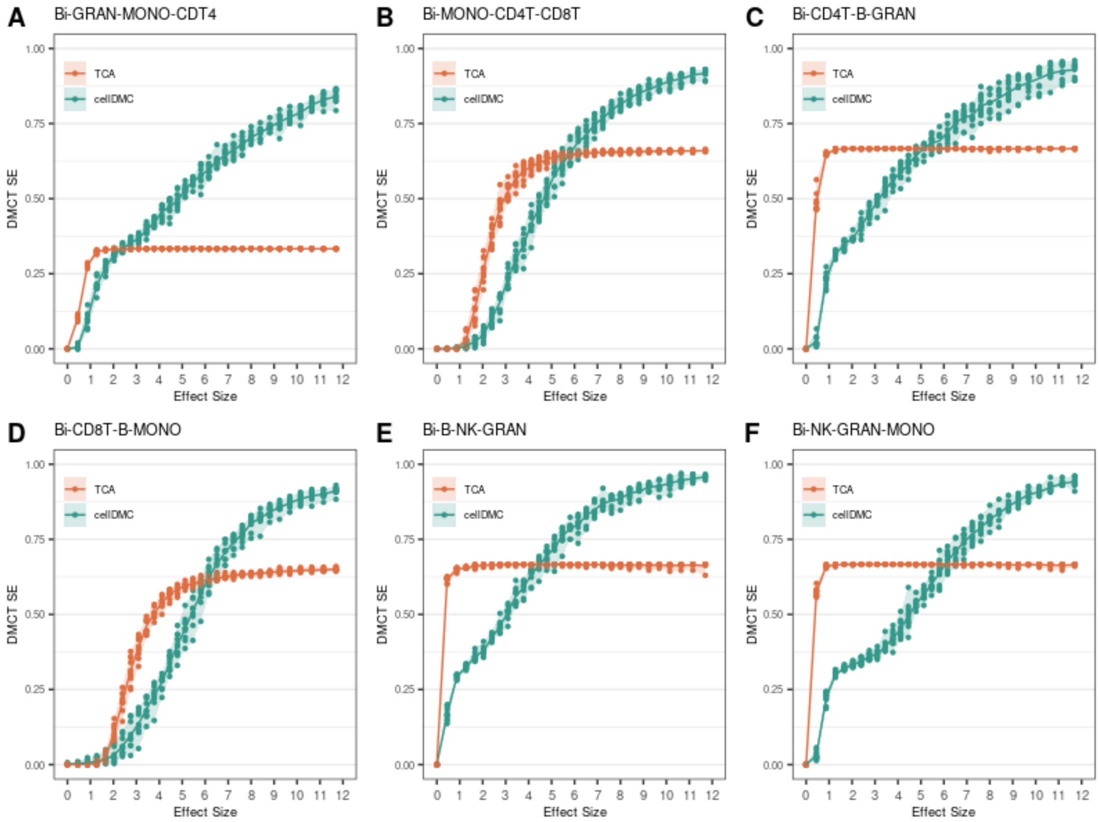


**Fig.S5: Comparison of the sensitivity (SE) in the bidirectional three altered cell-type scenario.** Plots of the sensitivity (SE) to detect DMCTs of CellDMC and TCA vs. the effect size for the scenario of bidirectional differentially methylation in 3 immune cell subtypes, as indicated above each plot. The mixtures are made up of a total of 6 immune cell subtypes. Results are for 10 Monte Carlo runs, where in each run 200 samples were simulated (100 controls and 100 cases). The solid lines show the mean sensitivity change over the 10 Monte-Carlo runs.

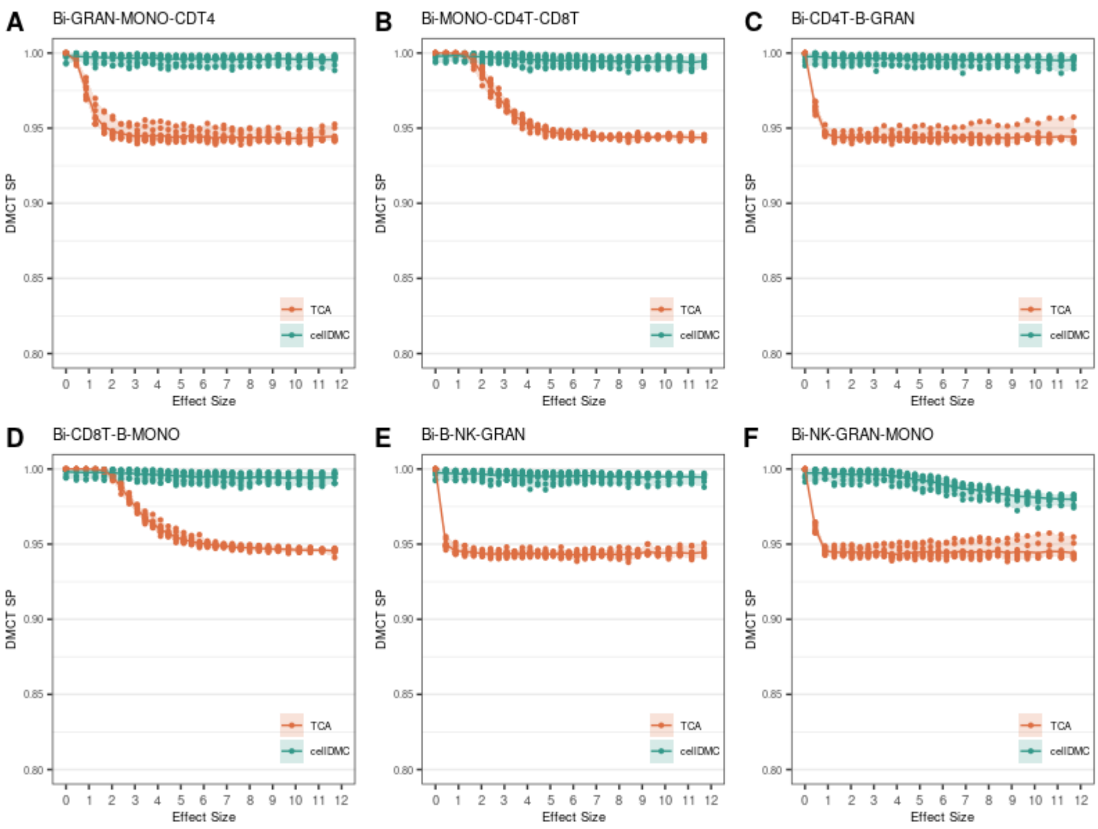


**Fig.S6: Comparison of the specificity (SP) in the bidirectional three altered cell-type scenario.** Plots of the specificity (SP) to detect DMCTs of CellDMC and TCA vs. the effect size for the scenario of bidirectional differentially methylation in 3 immune cell subtypes, as indicated above each plot. The mixtures are made up of a total of 6 immune cell subtypes. Results are for 10 Monte Carlo runs, where in each run 200 samples were simulated (100 controls and 100 cases). The solid lines show the mean sensitivity change over the 10 Monte-Carlo runs.

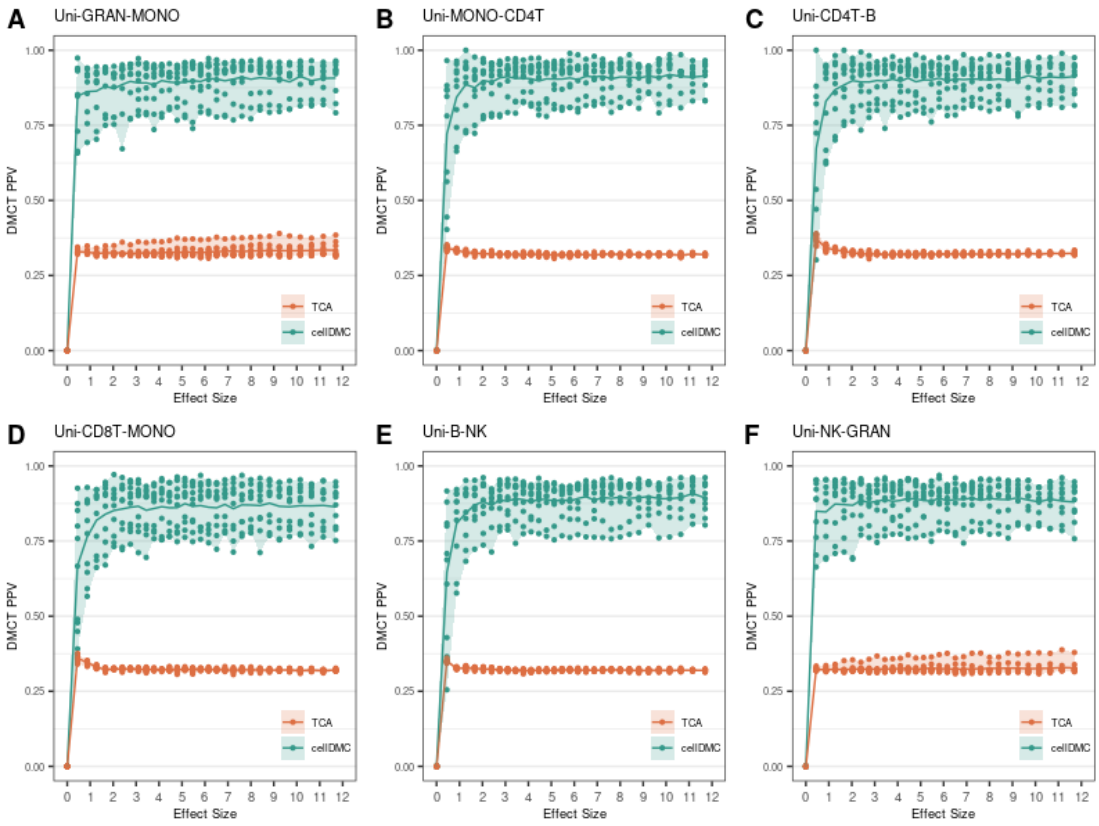


**Fig.S7: Comparison of the precision or positive predictive value (PPV) in the unidirectional two altered cell-type scenario.** Plots of the positive predictive value (PPV) to detect DMCTs of CellDMC and TCA vs. the effect size for the scenario of unidirectional differentially methylation in 2 immune cell subtypes, as indicated above each plot. The mixtures are made up of a total of 6 immune cell subtypes. Results are for 10 Monte Carlo runs, where in each run 200 samples were simulated (100 controls and 100 cases). The solid lines show the mean sensitivity change over the 10 Monte-Carlo runs.

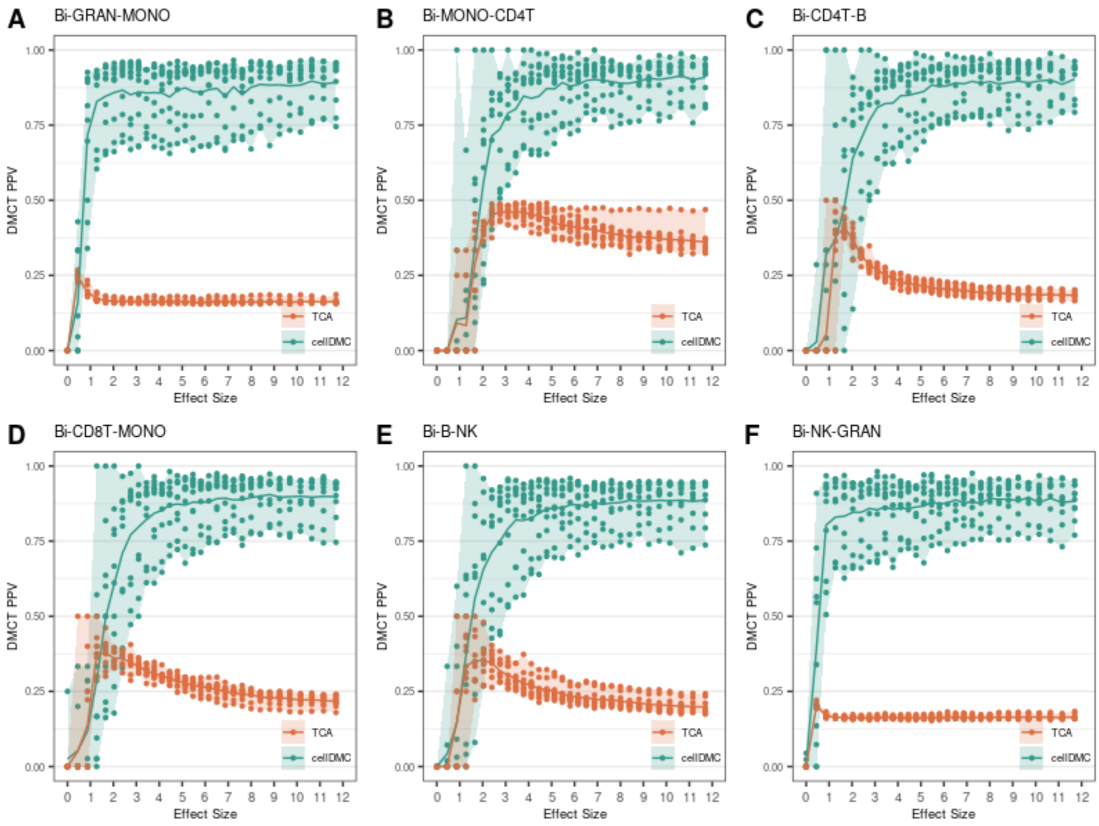


**Fig.S8: Comparison of the precision or positive predictive value (PPV) in the bidirectional two altered cell-type scenario .** Plots of the positive predictive value (PPV) to detect DMCTs of CellDMC and TCA vs. the effect size for the scenario of bidirectional differentially methylation in 2 immune cell subtypes, as indicated above each plot. The mixtures are made up of a total of 6 immune cell subtypes. Results are for 10 Monte Carlo runs, where in each run 200 samples were simulated (100 controls and 100 cases). The solid lines show the mean sensitivity change over the 10 Monte-Carlo runs.

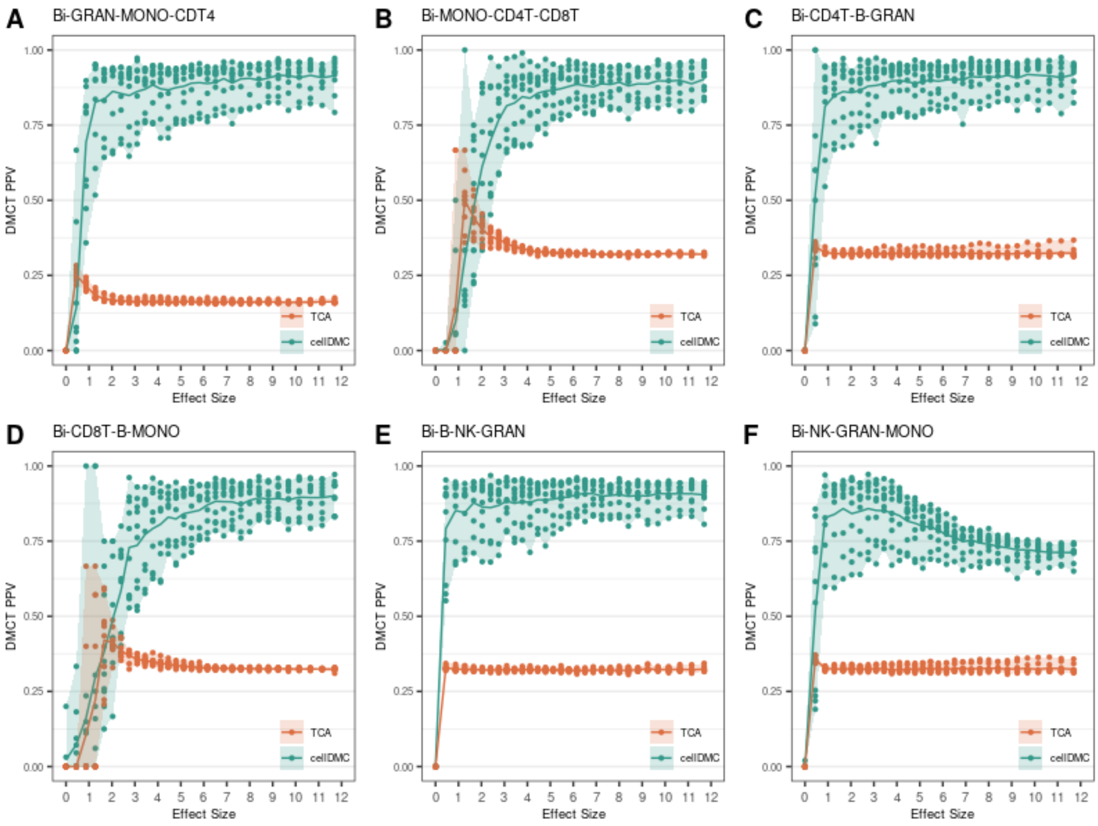


**Fig.S9: Comparison of the precision or positive predictive value (PPV) in the bidirectional three altered cell-type scenario .** Plots of the positive predictive value (PPV) to detect DMCTs of CellDMC and TCA vs. the effect size for the scenario of bidirectional differentially methylation in 3 immune cell subtypes, as indicated above each plot. The mixtures are made up of a total of 6 immune cell subtypes. Results are for 10 Monte Carlo runs, where in each run 200 samples were simulated (100 controls and 100 cases). The solid lines show the mean sensitivity change over the 10 Monte-Carlo runs.

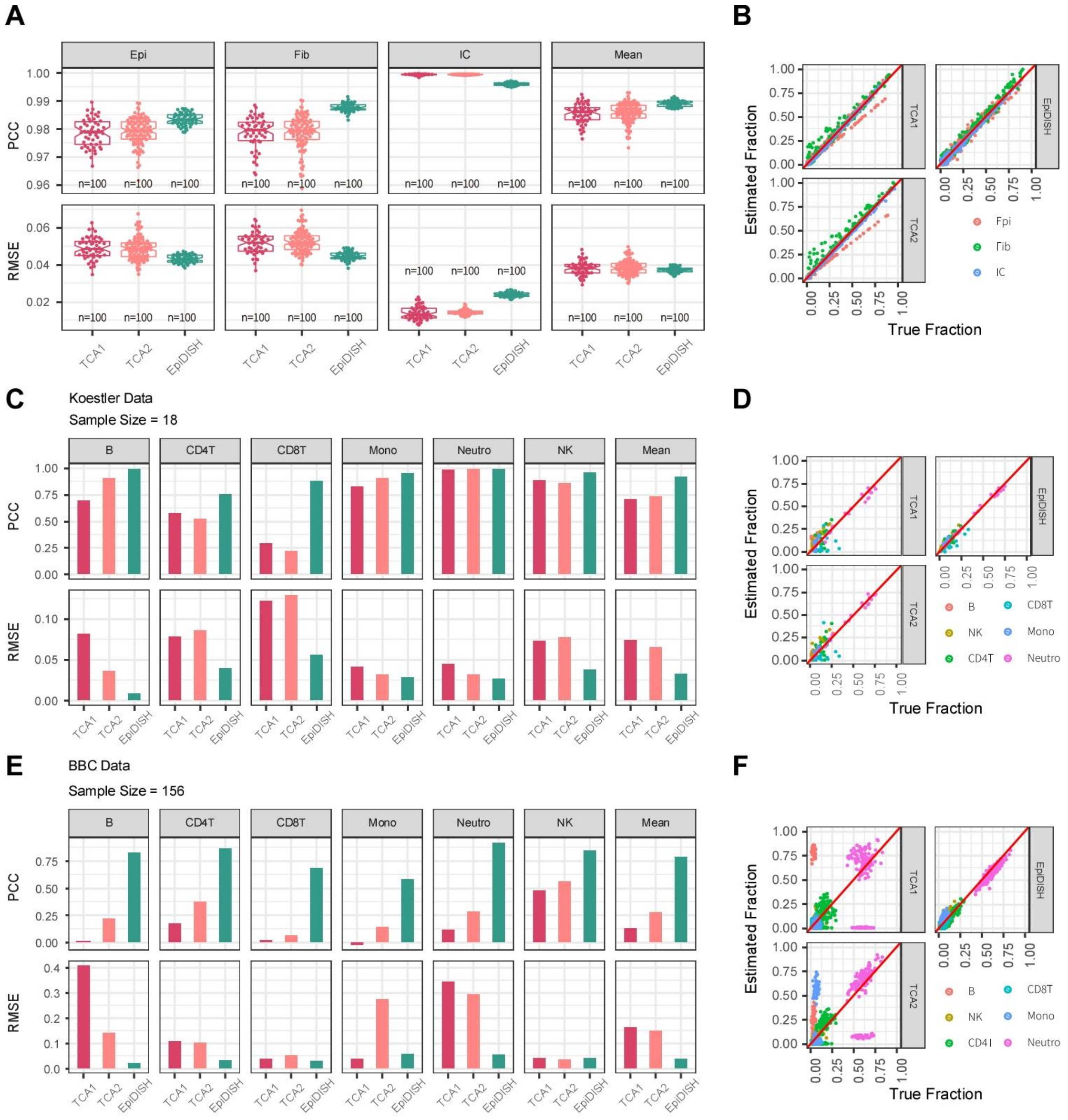

**Fig.S10: Evaluation of cell-type fraction estimates using TCA. A)** Boxplots of the Pearson Correlation Coefficient (PCC) and Root Mean Square Error (RMSE) between the estimated cell-type fraction and the true fraction for 3 cell-types (Epi=epithelial, Fib=fibroblast, IC=immune cell), for the case of 100 in-silico generated mixtures of epithelial (Epi), fibroblast (Fib) and immune cell (IC) subtypes. The last panel shows the mean PCC and RMSE over the 3 cell-types. Shown are the PCC and RMSE for 3 different methods of estimating the cell-type fractions: TCA1=using final TCA estimates initializing with EpiDISH, TCA2=using final TCA estimates initializing with EpiDISH plus noise, EpiDISH=cell-type fraction estimates derived using the robust partial correlation framework of EpiDISH [1]. **B)** Corresponding scatterplots of estimated vs. true cell-type fractions for the 3 cell-types and for each of the 3 different methods. **C-D)** As A-B), but now comparing the estimated cell-type fractions to the true fractions in 18 whole blood samples from Koestler et al [2]. In this dataset, the true fractions refer to either FACS cell counts or to known proportions for the experimentally derived mixtures, as described in Koestler et al [2]. **E-F)** As C-D), but now for a whole blood data set of 162 samples with matched FACS cell counts.

### Supplementary Methods

*Simulation model:* Our simulation models are based on the Illumina 450k DNA methylation (DNAm) profiles for 6 major immune cell subtypes (CD4+ and CD8+ T-cells, Granulocytes, monocytes, CD19+ B-cells and natural killer (NK) cells) from Reinius et al [3]. In this DNAm set there are 6 purified samples for each cell-type. In order to more realistically simulate real mixtures of these 6 cell-types in large numbers of samples, we first estimated means and standard deviations at each CpG site across each of the 6 cell types assuming an underlying beta-distribution using the method of moments. For generating the in-silico mixtures and for computational efficiency, we constructed the DNAm data matrices over a total of 1333 CpGs, made up of 1000 CpGs picked at random from the 450k array and the additional set of 333 "reference" CpGs which make our EpiDISH DNAm reference matrix [1]. These 333 reference CpGs are used to infer cell-type fractions in mixtures. A total of 200 in-silico mixtures were generated, i.e. 100 controls and 100 cases where DMCTs are defined. To define differentially methylated cell-types (DMCTs), that is differentially methylated cytosines within specific cell-types, we first selected at random 100 CpGs from the 1000 non-reference CpGs, but ensuring that these 100 CpGs exhibit either low (<0.2) or high (>0.8) average DNAm levels in the specific cell-types to be altered. This was done to ensure that we could consider a wide-range of effect sizes without having to worry about the bounded nature of beta-values. Thus, these levels of methylation represent the baseline levels in control samples, and thus for CpGs exhibiting low methylation in controls we would only consider hypermethylation in cases, and vice-versa for CpGs highly methylated in controls, we would only consider hypomethylation. In total, we considered 4 different DMCT scenarios: unidirectional change in one cell type, unidirectional changes in two cell types, bidirectional changes in two cell types and bidirectional changes in three cell types. Thus, in the unidirectional two cell-type scenario, a given CpG that is normally unmethylated in say granulocytes and B-cells, would exhibit hypermethylation in both cell-types in cases. Similarly, in the bidirectional two cell-type scenario, a CpG that is normally unmethylated in granulocytes and fully methylated in B-cells, would exhibit hyper and hypomethylation in granulocytes and B-cells, respectively. For each scenario, we considered a total of 60 different effect sizes with values ranging from 0 to approximately 12, with corresponding absolute DNA methylation differences in individual cell types (between cases and controls) varying from 0 to 0.6 maximally. Thus, these range of effect sizes capture effectively any realistic scenario. The calculation of effect size we use is given by the following equation:

$$effect\_size = \frac{\Delta\mu}{\sqrt{\frac{\sigma_1^2 + \sigma_2^2}{2}}}$$

where $\Delta\mu = |\mu_1 - \mu_2|$ is the absolute mean DNA methylation shift between case and control group, $\mu_1$ and $\mu_2$ are the mean DNA methylation levels in case and control group respectively, and where $\sigma_1$ and $\sigma_2$ are the variance of DNA methylation levels in case and control groups, respectively. Here we fixed $\sigma_1 = \sqrt{\frac{\mu_1(1-\mu_1)}{\mu_2(1-\mu_2)}} \cdot \sigma_2$ to ensure that the variance changes with the mean according to a beta-valued distribution.

Cell-type fractions to use for each in-silico mixture were obtained by sampling from a true immune cell subtype fraction distribution, as derived using EpiDISH on the Hannum et al whole blood dataset [4], which contains 656 Illumina 450k whole blood samples. Finally, we composed bulk methylation level at each CpG site by taking a linear combination of the simulated cell-type-specific levels across the 6 major immune cell subtypes, weighted by the corresponding cell type fractions. For every scenario and effect size, we ran 20 Monte-Carlo simulations.

*Implementation of CellDMC and TCA:* We implemented CellDMC [5] using the corresponding R package named "EpiDISH". CellDMC was run with the estimated cell type fractions from EpiDISH, which effectively corresponds to the true fractions, as shown by us previously [1, 5]. Case/control status was encoded as a binary phenotype. Correction for multiple-testing was done by estimating the FDR and calling significance at an FDR threshold of 0.05, which results in an output matrix containing predicted DMCT(s) per cell-type, as well as their direction of DNAm change.

We applied TCA using the corresponding R package called "TCA" [6]. We ran TCA with initial cell-type fraction estimates as given by EpiDISH, that is we use the same estimates as when running CellDMC. In our simulation model, there are no other sources of variation besides phenotype and cell-type, so the parameters C1 and C2 were set to NULL. From the TCA-fit, we subsequently inferred the data-tensor defined over cell-types, CpGs and samples. Thus, for each cell-type, this data tensor defined a corresponding cell-type specific DNAm data matrix, on which we subsequently performed an ordinary linear regression to derive t-statistics and P-values of association against case/control status. Multiple-testing correction was performed as before using an FDR < 0.05 threshold, resulted in an output matrix containing predicted DMCT(s) per cell-type, as well as their directionality of DNAm change.

*Definition of sensitivity (SE), specificity (SP) and positive predictive value (PPV):* All these measures were first defined for each cell-type separately, and subsequently, values were averaged over all relevant cell-types. For a given cell-type that was chosen to be altered, the sensitivity to detect DMCTs in this cell-type was defined as the ratio of the predicted true number of DMCTs divided by the total number of true DMCTs. Correspondingly, the specificity for this cell-type was defined as 1- FPR, where the false positive rate (FPR) is the ratio of false positives to the total number of true negatives. Of note, a true DMCT predicted to be a DMCT but where the predicted directionality of DNAm change is wrong, was counted as a false positive. Likewise, a true negative CpG predicted to be a DMCT (regardless of directionality) also defines a false positive. Finally, the precision or PPV for a given cell-type was defined as the fraction of correctly predicted DMCTs (with correct directionality of change) among all CpGs declared to be DMCTs.

*Real DNAm datasets with matched FACS cell counts:* We used the Illumina 450k dataset from Koestler et al [2] consisting of 18 samples, of which 6 were whole blood (WB) and 12 were experimentally reconstructed "whole blood" mixtures. This dataset is available from GEO under accession number GSE77797. In the case of the 6 whole blood samples, flow-cytometric cell count estimates for the 6 major blood cell subtypes were available. For the experimental mixtures, the mixing proportions were determined by the experimentalist and therefore known without error. DNAm data was normalized and processed as previously described [1]. In addition we analysed Illumina EPIC DNAm data for a total of 162 whole blood samples with matched FACS cell counts. The EPIC DNAm data was processed using minfi [7, 8] and BMIQ [8]. This dataset is available from https://www.biosino.org/node/index under accession number XXX (to be determined).
